# Kin structure and roost fidelity in greater noctule bats

**DOI:** 10.1101/675215

**Authors:** João D. Santos, Christoph F.J. Meyer, Carlos Ibáñez, Ana G. Popa-Lisseanu, Javier Juste

**Author notes:** Correspondent: João Santos, present address: Cirad, AGAP, TA A-108 / 03, Avenue Agropolis, 34398 Montpellier Cedex 5, France.

## Abstract

Roost fidelity is an important aspect of mammalian biology. Studying the mechanisms underlying philopatry can help us understand a species’ energetic requirements, ecological constraints and social organisation. Temperate bat species notably exhibit a comparatively high degree of female philopatry, resulting in maternity colonies segregated at the mitochondrial level. We focus on the greater noctule, *Nyctalus lasiopterus*, to study this behaviour in more depth. We make use of microsatellite data for 11 markers across 84 individuals residing in Maria Luisa Park in Seville, Spain. At the time of sampling, this urban park boasted the highest number of registered bats of this species, among which three social groups were observed to segregate spatially. We studied the distribution of pairs of individuals across filial relationship categories and relatedness estimates relative to the social group of each individual. This analysis was complemented by information on roost-use frequency among a subset of genotyped bats. We found no significant relationship between roost use and genetic distance, but there was evidence of higher group-sharing with increased levels of relatedness. Mother-daughter pairs shared the same group more often than expected, as did pairs of individuals of relatedness above 0.36. We discuss the implications of these results in terms of the behavioural ecology of temperate bats and for conservation efforts aimed at preserving them.

## INTRODUCTION

Female philopatry, or fidelity to a birth site or region, is a common trait among mammals (Greenwood, 1980; Clutton-Brock & Lukas, 2012), including bats (Burland & Wilmer, 2001). Bats display an exceptionally high degree of female philopatry and colony fidelity (Kerth et al., 2008). While this can be partly explained by the temporal stability and sparse distribution of the roosts used by cave-dwelling species (Lewis, 1995; Kerth, 2008), it is still common in tree- and foliage-roosting bats, for which roost availability may not constitute a limiting factor (Kerth, 2008). Philopatry in female temperate tree-roosting bats has thus been attributed to familiarity with the natal area, roost and foraging locations, or the maintenance of social relationships (Kerth, 2008; Olivera-Hyde et al., 2019).

Sociality theory posits that philopatry and traits linked to colony fidelity were selected for in order to take advantage of the benefits of group living. Indeed, reported interactions among females of bat colonies include communal nursing (Wilkinson, 1992), social grooming (Wilkinson, 1986; Kozhurina, 1993; Kerth 2008), food sharing (Carter & Wilkinson, 2013) and information transfer about roost sites and foraging locations (Wilkinson & Boughman, 1998; Kerth & Reckardt, 2003; Chaverri et al., 2010). Strong female-biased philopatry together with high longevity in bats (Barclay & Harder, 2003) should lead to the formation of matrilineal societies and open the door for the fixation of cooperative behaviours through kin selection (Hamilton 1964; Emlen & Oring, 1977; Clutton-Brock, 2009). However, the extent to which these interactions are maintained by kin selection is unknown. For example, food sharing in *Desmodus rotundus* was reported to be more dependent on reciprocity than relatedness (Carter & Wilkinson, 2013). Likewise, information transfer in both the evening and Bechstein’s bats showed no evidence of kin selection (Wilkinson, 1992; Kerth & Reckardt, 2003). For other behaviours, such as association and grooming, empirical support for different species is incongruent (Metheny et al., 2008; Patriquin et al., 2013).

Studies monitoring roost associations have shown females will roost preferentially with some females rather than others, resulting in a non-random association pattern within the colony (Willis & Brigham, 2004; Metheny et al., 2008; Patriquin et al., 2013; Nado et al., 2017). Because tree-roosting bats occupy hollows not large enough for more than a small number of individuals, regular fission and fusion of subgroups could help maintain larger social groups spread over a given area (O’Donnell, 2000; Patriquin et al., 2016). A recent review of studies reporting pairwise association and genetic relatedness data across nine species of bats spanning four families found no correlation between the strength of associations between kin and the occurrence of complex cooperative behaviours (Wilkinson et al., 2019). Rather, the authors observed that species that switched roosts often tended to present stronger associations between relatives. Understanding the causes underlying philopatry in this context is rendered particularly difficult. The kind of philopatry at play – to the natal area (which may not necessarily relate to social behaviour) or to the natal colony, becomes elusive.

The greater noctule, *Nyctalus lasiopterus* (Schreber, 1780), is one of the largest and rarest European vespertilionid bats (Ibáñez et al., 2004). Greater noctules typically roost in large, old trees (Alcalde et al., 2016). The sparse number of existing records suggests that the species’ distribution is circum-Mediterranean and patchy from Morocco to Uzbekistan (Ibáñez et al., 2004). Western Andalusia, Spain, is unique for harboring several known maternity colonies of greater noctules in three main breeding areas (Santos et al., 2016). Females consolidate colonies in spring to breed, and remain until late August, when they disaggregate and disperse, presumably to mating and hibernation sites (Ibáñez et al., 2009), although some cases of permanent residency have been observed (C. Ibáñez and J. Juste, unpublished data). Few males have been found in these colonies, contrasting with other sites in northern Spain and Europe (Ibáñez et al., 2009).

In 2008, a study on the social structure of the greater noctule population of Maria Luisa Park (MLP) in Seville (Spain) revealed it to follow a fission-fusion structure, with bats switching roosts on average every three days and segregated in three spatially distinct social groups (9-17% overlap), whereby the same tree holes were rarely used by bats of different groups (Popa-Lisseanu et al., 2008; Fortuna et al., 2009). Moreover, the foraging areas of the three groups (extending over 40 km away from the park) overlapped almost completely. Popa-Lisseanu et al. (2008) further reported that females were loyal to their groups for at least five years, and that juvenile females returned to their natal roosting areas during the study period. The authors concluded that the three groups observed within the park could be considered independent maternity colonies, defined as groups of reproductive females roosting together. However, more recently, a genetic analysis encompassing the region of Andalusia, Spain, and including three maternity colonies apart from Seville (which was considered as a single population), found significant mitochondrial structure at the regional scale, but not within the park (Santos et al., 2016).

In this study, we narrowed the geographic focus of our previous work on regional genetic structure (Santos et al., 2016) to the urban park in Seville. Using bi-parentally inherited markers (microsatellites), we asked whether the status of colony attributed to the different social groups within the park could be verified genetically. If the groups represent socially and demographically separated colonies, then we expect this behaviour to lead to genetic isolation, detectable as: i) higher within-than between-group relatedness, conditional on the age of each group, since higher within-group relatedness was not observed during colonization (Santos et al., 2016), ii) a similar deviation in the distribution of related pairs of females across groups to that reported by Santos et al. (2016) across the region (throughout the article, we refer to the groups within the park as groups or social groups, and to the Seville population as population or colony). Furthermore, within our genetic dataset, we include samples from individuals used by Popa-Lisseanu et al. (2008) to identify temporal roosting associations. Because the three groups are both physically and socially defined, we used this to refine our initial hypothesis: H0) group identity, physical or social, does not impact individual decisions to return – genetic structure should be absent, genetic and roosting data disassociated; H1) Group identity alone has inherent impact – higher within-group relatedness should be observed but no roosting/genetic structure; H2) Group identity impacts dyad structure because *N. lasiopterus* females chose to roost with relatives – both dyad structure and roosting/genetic association should be observed. H3, the absence of genetic isolation in the presence of roosting/genetic association, was not seriously considered as it would imply a more complicated scenario of familial interactions. Regarding our understanding of the philopatric behaviour of female giant noctules, H0 corresponds to a coarse site fidelity, and could be explained by the limited number of alternative roosting opportunities in the region. H1 implies a more specific preference for certain roosts or individuals, and H2 would indicate the maintenance of kin-relationships as the basis for the return.

## Materials and Methods

### Focal Species and Study Population

Very few breeding colonies of *N. lasiopterus* are known outside of the Iberian Peninsula: France, Hungary, Slovakia, Belarus, Ukraine and Russia (Uhrin et al., 2006; Estók & Gombkötő, 2007; Dubourg-Savage et al., 2013; Dombrovski et al., 2016; Vlaschenko et al., 2016; Kovalov et al., 2018).

The study population is located in MLP, Seville (37°22′29″N, 5°59′19″W). The 122-year-old park extends over 38 ha and contains many large old trees that bear enough cavities to harbour the three distinct colonies. Most roost trees belong to *Platanus* spp., *Gleditsia triacanthos, Sophora japonica* and *Washingtonia filifera*. With a population previously estimated at ca. 500 individuals, but now in dramatic decline due to an invasive parakeet species (Hernández-Brito et al., 2018) that is taking over the roosting holes, it was once the largest known breeding site of greater noctules (Popa-Lisseanu et al., 2008).

### Sampling

We analysed biopsy samples of 82 adult female bats from MLP collected from 2004 to 2007. Each bat was assigned to one of three areas of the park identified by Popa-Lisseanu et al. (2008) as characteristic of different social groups. Assignment was based on the tree in which individuals roosted in when captured. Group designations (I, II and III) were maintained from that earlier study. Twenty-four individuals were sampled from each group. Samples included 25 individuals that were radio-tracked to study the roost-switching behaviour of the population that was used to identify and define the social group dynamics in the previous study (Popa-Lisseanu et al., 2008).

Capture and experiments were performed under permit of the Ethical Committee of EBD-CSIC and the techniques used meet the guidelines published by the American Society of Mammalogists (Gannon and Sikes, 2007) on the use of wild mammals in research.

### Molecular Methods

Total DNA was extracted from 3 mm diameter wing punches using a modified version of the salt-based protocol developed by Aljanabi & Martinez (1997).

All individuals were genotyped at 11 nuclear microsatellite loci: Nle 2, 3 and 6-11 (developed for *Nyctalus leisleri*, see Boston et al., 2008); EF4 (developed for *Eptesicus fuscus*, Vonhof et al., 2002); P20 and P217 (developed for *Pipistrellus sp*. Kaňuch et al., 2007). All were tested in muscle tissue prior to genotyping (see Supplementary Methods for a detailed description of DNA extraction, amplification, sequencing, and microsatellite genotyping) and successfully employed in our previous study (Santos et al., 2016). Labelling followed Schuelke’s procedure (2000).

### Data Analysis

All nuclear markers were in linkage equilibrium as assessed with FSTAT v. 2.9.3.2 (Goudet et al., 1995). Using CERVUS, we estimated observed and expected heterozygosities, as well as deviations from Hardy-Weinberg equilibrium (HWE). Of the 11 microsatellites, three (Nle6, P20 and P217) deviated significantly from HWE (see Supplementary Table S1).

Pairwise and mean relatedness values (*r*) were estimated using ML-Relate (Kalinowski et al., 2006), which implements a corrected maximum-likelihood method for estimating relatedness that allows loci with null alleles to be incorporated into the analysis (Wagner et al., 2006). Three of the 11 loci used in this study showed the highest frequencies of null alleles (min=0.097, mean=0.187). Incidentally, this might have been the cause of the departure from HWE, as it would simulate a heterozygote deficiency. Because we did not want to abdicate the inferential power of three loci, we resorted to the method by Wagner et al. (2006), implemented in ML-Relate, and kept all 11 loci. Using this software, pairwise relations were classed as Parent-Offspring (PO), Full-Sibling (FS), Half-Sibling (HS), or Unrelated (U), and each attributed a value *r* of pairwise relatedness. Principal component analysis (PCA) was performed on the squared matrix of pairwise relatedness values.

With reference to the three groups within the park, pairwise relationships were classed according to whether or not both individuals had been captured within the same group. We tested the null hypothesis that each group shared the same population mean as the whole population. Firstly, analysis of molecular variance (AMOVA) was conducted between global and within-group data sets of relatedness estimates, using the function ‘f_oneway’ of the python package ‘scipy.stats’ (Oliphant, 2007). We then compared within-group relatedness to relatedness across the park independently for each group using a randomization test. We relied on two-tailed t-tests as implemented in the function ‘ttest_ind’ of the python package ‘scipy.stats’ (Oliphant, 2007). We approximated the distribution of the test statistic by comparing relatedness estimates across the park to group-sized samples of randomly selected relatedness values. The procedure was repeated 1000 times for each group. The final *p*-value was calculated as the proportion of sampled statistics below that observed for each group.

Regarding group choice, we assumed as H0 that the probability of choosing the same group as another individual at random follows a binomial distribution. We further considered this probability to be conditional on the relative size of the group, not the relative sample size. Then, the probability P(O) of pairs of individuals falling across groups was estimated as P(O) P_I_(1 − P_I_) + P_II_ (1 − P_II_) + P_III_(1 − P_III_). The values P_I_, P_II_ and P_III_were estimated as the proportions of individuals registered to roost in groups I, II and III at the time of sampling (81, 61 and 114 bats respectively, Popa-Lisseanu et al. 2008), so that P(O) = 0.644. The probability P(I) of two individuals choosing the same group was estimated as 1 – P(O) = 0.356. We compared the expected and observed proportions across the data set and within the classes of relation PO, FS, HS and U, and then studied the deviation of observed within-group pairs as a function of relatedness values. We estimated the upper-tail *p*-value of observed proportions of same-group pairs within each class assuming a binomial probability *P*(*I*). We further estimated upper-tail *p*-values of the proportion of dyads falling within the same group along windows of relatedness. Proportions were calculated among relationships presenting values within a 0.15 window of relatedness around a central value ranging between 0.075 and 0.5 in increments of 0.01. Window size was selected based on the minimum, median and standard deviation of the number of relationships across windows of a given size (see Supplementary Figure S1).

To explore the relationship between genetic relatedness and roost use we resorted to the matrix of pairwise roost use similarity among 15 bats radio-tracked in 2004 used by Popa-Lisseanu et al. (2008) to infer the existence of social groups within the park (Supplementary Figure S2). Similarity was estimated using the Freeman-Tukey statistic (Krebs, 1989). All individuals considered in that study were genotyped. A Mantel test of the two distance matrices was performed using the mantel.rtest function of R library ade4 (Dray and Dufour 2007) with the default number of repeats.

We constructed network graphs of roosting associations and parent-offspring dyads. Roosting association edges were created as dyads presenting a similarity value above 0.05 (Freeman-Tukey statistic). Networks were constructed using the Fruchterman-Reingold algorithm as implemented in the Networkx python package (Aric et al., 2008).

## Results

We first evaluated genetic structure across the park considering its known subdivisions. No genetic structure was observed through PCA (Fig. 1A), and one-way AMOVA of within-group relatedness estimates was non-significant (*p* = 0.16, d.f. = 1133). We then considered the interplay of group assignment, genetic relatedness and relationships. Of the total number of possible associations within the park and groups (every female paired with every other female), the percentage of those exceeding a relatedness degree of 0.25 accounted for 6% of the total (Table 1). However, around 81% of females shared their group with at least one close relative (*r* > 0.25, Table 1), while 95% shared the park (considering close relatives found in both the same and different groups). We focused on each group to ascertain whether mean relatedness exceeded expectations under randomness. The result of the t-test of pairwise relatedness means between total and group III dyads was significant (p < 0.01, Fig. 1B), as was that between total and group I (p < 0.05, Figure 1B).

**Table 1:** Mean relatedness estimates within groups and across the whole colony in Maria Luisa Park, Seville, percentage of associations with *r* > 0.25 (close relatives) among total possible associations, and percentage of females with close relatives within their groups as well as the whole colony. The total sample consists of 84 individuals, 24 from each group, resulting in a total of 3486 possible dyads.

| Group | % associations |  |  |  |
| --- | --- | --- | --- | --- |
|  | *Mean r* | *SD* | *r>0.25* | % |
| I. | 0.056 | ±0.106 | 6.3 | 89.2 |
| II. | 0.050 | ±0.099 | 5.6 | 75.0 |
| III. | 0.051 | ±0.099 | 5.6 | 78.6 |
| **Colony** | **0.059** | **±0.098** | **5.7** | **95.2** |

**Fig. 1.**
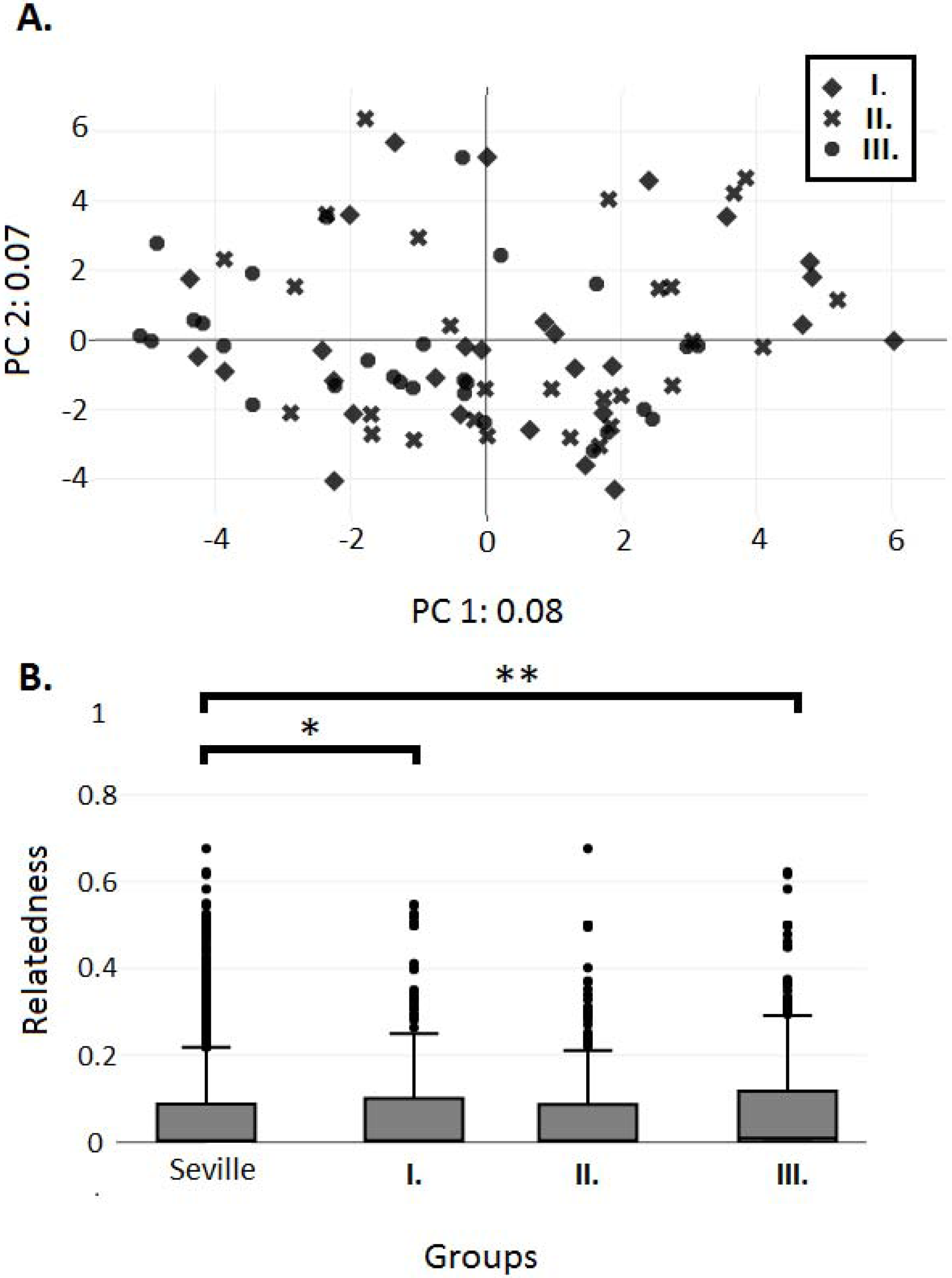
Pairwise relatedness between adult female *Nyctalus lasiopterus* within and among social groups in Maria Luisa Park, Sevilla, Spain. **A**) Biplot of a principal component analysis based on a relatedness matrix. **B**) Boxplot of pairwise relatedness across the pooled data set and within each social group. t-tests were calculated for group and colony sample means for each of the three groups (*p*-values shown above).

We also looked at the impact of group-sharing on relationship categories and relatedness estimates. The proportion of parent-offspring pairs that shared the same group differed significantly (*p* < 0.001) from random expectations (Fig. 3). The same proportions for FS, HS and U pairs did not deviate significantly from the expected proportions (Fig. 2). However, a decreasing trend is observed when these groups are considered in order of decreasing relatedness.

**Fig. 2.**
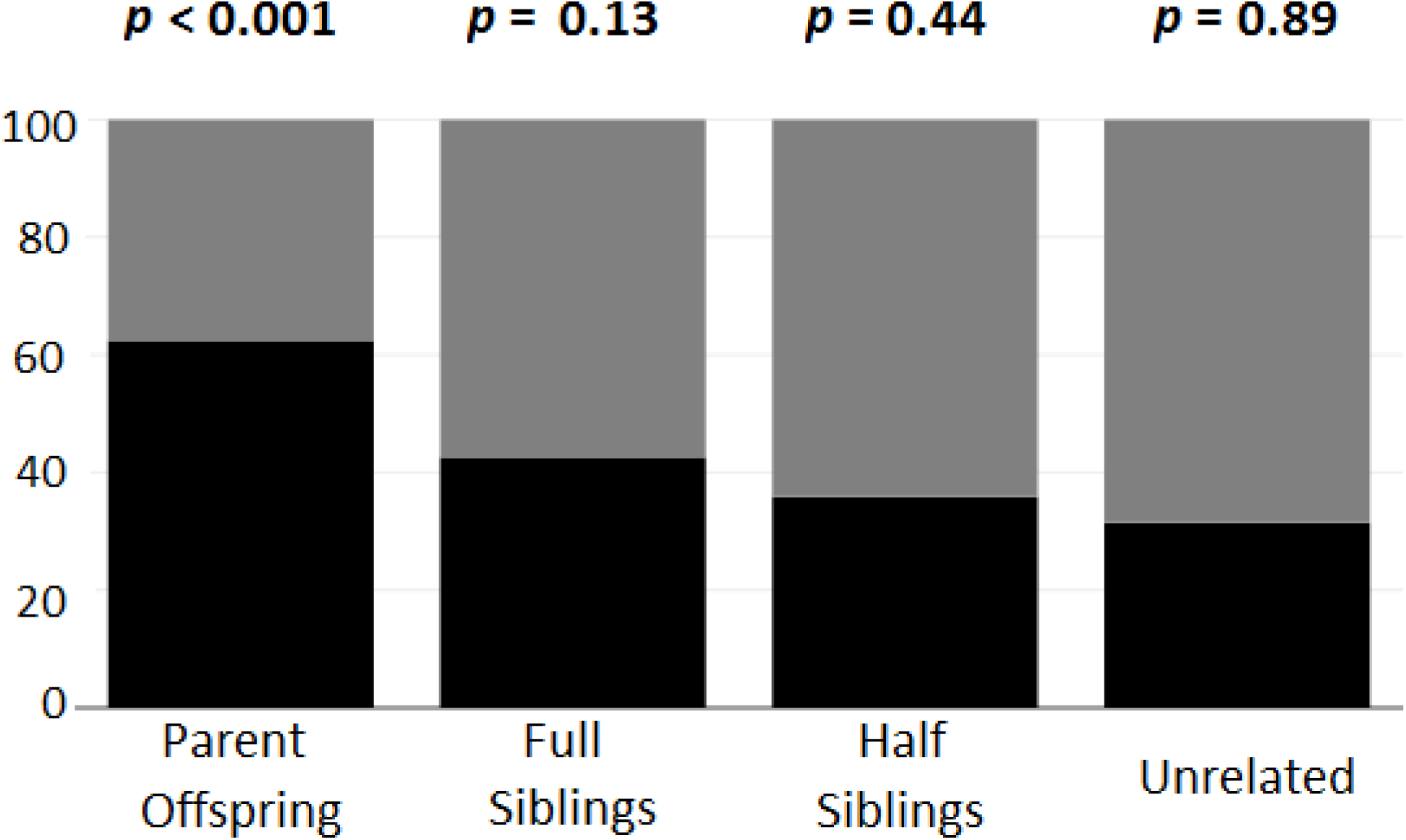
Percentage of pairwise relations within and among social groups across relatedness categories. Dyads were classed into relation groups using ML-Relate. Individual dyads were further classed according to whether they were caught within the same areas of Maria Luisa Park. Grey: proportions of dyads between individuals from different groups; black: proportion of dyads between individuals from the same group. *P*-value of observed proportion of pairs from the same group for each class shown in bold. Categories PO, FS, HS and U consisted of 40, 47, 441 and 2958 dyads respectively. *P*-values were derived assuming a binomial probability of two individuals sharing the same group *P*(*I*) = 0.356.

**Fig. 3.**
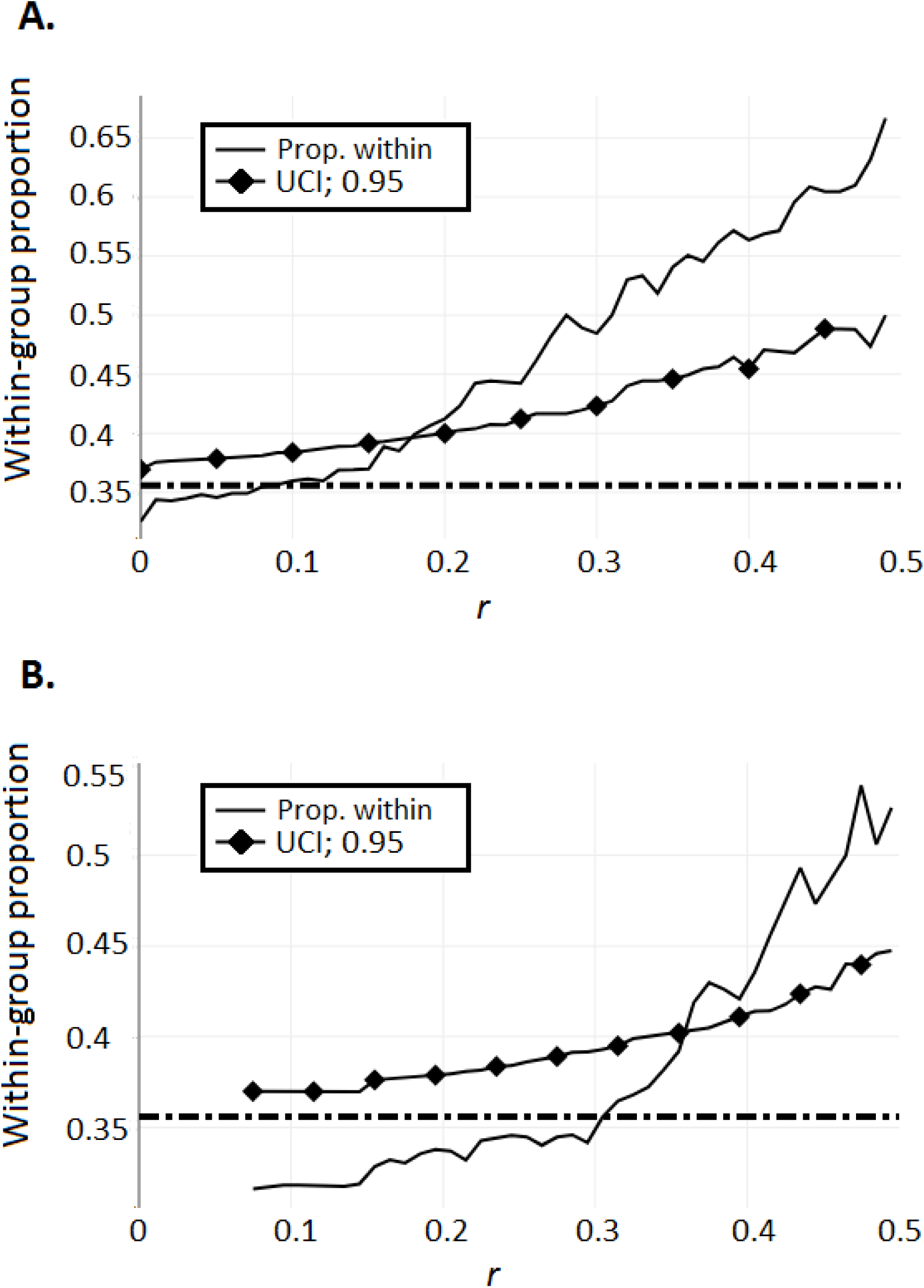
Proportion of group sharing as a function of pairwise relatedness. Pairs of individuals were classified according to whether both were captured in the same area of Maria Luisa Park. This proportion was then studied in relation to the estimated relatedness of each pair. **A**) Proportion of dyads within the same group with relatedness values above abscissae. **B**) Proportion of dyads within relatedness windows that fall within the same group (window size= 0.15 *r*, step= 0.03 *r*). The upper-bound 95% confidence interval (UCI, lozenge-line) was estimated from a binomial distribution of *p* = (*I*). The dashed line indicates the expected proportion *P*(*I*) of within-group pairs.

When the same test is performed on pairs of individuals of increasing relatedness, the proportion of these pairs falling within the same group rapidly becomes significant (r > 0.18, Fig. 3A). When only pairs within a moving window of relatedness are considered (size= 0.15, step= 0.01), the same proportion becomes significant above a relatedness level of 0.36 (Fig. 3B).

Finally, we assessed the effect of roosting association on genetic relatedness. A Mantel test revealed no significant association between the two variables (r = 0.06, simulated *p*-value = 0.27). This subset of individuals represented 37 half-sibling relationships, 2 full-sibs, 1 parent-offspring pair and 65 unrelated dyads. Roost association between the mother-daughter pairs was above average, but not significantly so (P = 0.09).

## Discussion

Female fidelity to breeding colonies has been reported in a number of temperate bat species (Rossiter et al., 2002; Castella et al., 2008; Juste et al., 2009), leading, in the extreme, to the formation of closed societies as in *Myotis bechsteinii* (Kerth et al., 2000). The existence of among-colony mitochondrial structure at the regional level supports a certain degree of philopatry among female *N. lasiopterus* (Santos et al., 2016). Our results confirm that this behaviour extends to social groups within the population of Seville. This behaviour leads to a proportion of female pairs sharing the same group that increases with relatedness, becoming significant at higher levels (r > 0.36) and resulting in a substantial proportion (62.5%, 25 / 40) of predicted mother-daughter pairs sharing the same social group. While this value is below that observed for the strictly philopatric Bechstein’s bat (Kerth et al., 2002), it is considerably higher than reported in big brown bats (9%; Vonhof et al., 2008). We further find that two of the groups present significantly higher relatedness levels than the park as a whole, indicating that this behaviour is potentially stable. The fact that group II females do not exhibit higher average genetic relatedness could be explained if this group was recent, since group formation in this species is not dependent on kin (Santos et al., 2016).

The reasons for philopatric behaviour have been widely discussed (Moussy et al., 2013), and generally attributed to the females’ acquaintance with their natal area, to facilitate access to foraging and roosting sites, and to the benefits they might derive from social relationships and cooperative behaviours with conspecifics (Kerth, 2008; Kerth et al., 2011). We tested for the effect of roosting associations, previously shown to follow the same park sub-sections, and found a lack of correlation between roost use and genetic relatedness. Despite the limited power of this analysis due to small sample sizes, our results thus seem to refute our initial hypotheses H0 and H2 in favour of H1: females tend to return to natal roosting areas, but this choice is not driven by a desire to roost with kin. We can also discard explanations based on facilitated access; we are not aware of differences in roost characteristics between the three areas of the park that could account for the patterns observed, and the three groups have completely overlapping foraging areas (Popa-Lisseanu et al., 2009). Philopatric behaviour in Seville can thus be explained either by familiarity with the natal area, or the maintenance of social relationships that are not kin-based. Associative learning is the imprinting of roosts by non-volant young when being carried by their mothers between day roosts (O’Donnell, 2000). Based on reports of observational learning, O’Donnell (2000) reasoned that if young *Chalinolobus tuberculatus* could have their mother’s social group pool of favourite roost trees imprinted through repetitive roost switching, group structure would be maintained. The same mechanism could be at play in *N. lasiopterus*.

The alternative hypothesis is that roost fidelity functions to maintain social bonds, helping returning females reconnect with acquaintances made at specific roosts. Studying the social organization of a maternity colony of noctule bats *Nyctalus noctula* in captivity, Kozhurina (1993) reported the best predictors of social organization to be age and sex (92.6 % classification accuracy versus 29.6 % for family membership) and that the disparity between age classes remained over time. It is important to add that occasions of social grooming occurred strictly among members of the same social groups. A more recent study of the temporal and genetic components of associative dynamics in *Nyctalus leisleri* also found a stronger impact of cohort year on the composition of social groups inferred through network analysis (Nado et al., 2017). As pointed out by the authors, although genetic relatedness was also observed to significantly impact association between dyads, this could be the result of different genetic profiles between cohorts due to male-biased gene flow. Cooperative behaviours and long-term associations among colony members, sometimes spanning several years, have been described in numerous bat species (Wilkinson & Boughman, 1998; Carter & Wilkinson, 2013). Kozhurina (1993) further observed that noctule bats start forming associations in their first year. If this is true for greater noctules, maintaining these bonds, either for the comfort derived from familiarity and predictability (e.g. reduced aggression, reduced demands on limited attention capacity already strained by pup-rearing), or the increased likelihood of reciprocal cooperative behaviour, could explain why individuals tend to return to the area of the park they were born in. This theory has the advantage of providing an explanation for the formation of social groups, which are a pre-requisite for the associative learning hypothesis.

In summary, we found that female *N. lasiopterus* will seek to return to the specific areas or social groups they were born in, not just the region or breeding area, more often than expected by chance. We further found that roost associations are not kin-based. While these findings support the hypothesis that group structure is maintained through philopatry (Popa-Lisseanu et al., 2008), it is likely that neither associative learning nor sociality account for this behaviour entirely, but that instead both play a role. Both hypotheses explain the specificity of natal philopatry in greater noctules. Associative learning has the obvious adaptive advantage of allowing young adult females to benefit from their mothers’ successful choice of colony. Strong social bonds leading to cooperative behaviours, such as allo-nursing or information sharing, bring fitness benefits to nursing mothers with already high energy expenditures, an explanation which seems particularly plausible for females of this species in light of their challenging carnivorous habits during part of the year (Dondini and Vergari, 2000; Ibáñez et al., 2001; Popa-Lisseanu et al., 2007). While both hypotheses seem plausible, each rests on certain assumptions regarding roosting behaviour or social interactions that lack confirmation in this species. There is one case of reported co-operative nursing in captivity (Kozhurina, 1993), but no study to date has captured roosting associations or behaviours with enough precision to infer the correlates of social organization in maternity colonies of *N. lasiopterus*. Likewise, insufficient information is available on roost fidelity to assess how it correlates with breeding status or to verify roost reuse. Popa-Lisseanu et al. (2008) did report a decrease in roost switching during lactation, an observation that perhaps failed to reach significance due to lack of statistical power. Repeating this experiment with a larger sample size could be informative since a significant reduction in roost switching would render the associative learning hypothesis less likely. In the meantime, regardless of the mechanism involved, we argue that in light of our results, and those of previous studies in this park, it is safe to re-classify the social groups in MLP as separate colonies.

Determining the relative importance of associative learning and social bonds in this species is crucial, as it could direct future conservation efforts regarding the few known maternity colonies. In order to discriminate between these two hypotheses, future studies should focus on roost fidelity and roosting patterns as well as on long-term associations. Evidence of roost reuse by individual females, as well as varying roost fidelity according to breeding status would point to associative learning. These will have to be compared with the frequency of long-term associations within the same colonies in order to determine the relative importance of social bonds. More generally, discovering the mechanism behind philopatry represents an important step in our understanding of the thought processes behind the individual decisions that structure mammalian societies, and has implications for conservation policies aimed at preserving healthy populations.

## Acknowledgements

J. Nogueras and C. Ruiz helped collecting bat samples. We particularly thank J.L. García-Mudarra and J.M. Arroyos-Salas for valuable advice and their technical insight. We also acknowledge the staff of the Servicio de Parques y Jardines de Sevilla for their help and continuous support of our research. Logistical support was provided by the Laboratorio de Ecología Molecular, Estación Biológica de Doñana, CSIC (LEM-EBD). The regional government of Andalusia provided permits for collecting samples and handling of bats. This study was partially funded by the Spanish MICINN CGL 2009-12393, the PPNN 021/2002 and 1981/2010. JDS would like to acknowledge support from the Erasmus Student Mobility program.

## Supplementary Data

Supplementary Methods – Detailed description of DNA extraction, purification, sequencing and genotyping.

Supplementary Table S1 – Summary statistics and PCR specifications for each locus over all colonies.

Supplementary Figure S1 - Exploratory analysis of the impact of window size on sampling across relatedness steps.

Supplementary Figure S2 - Intra-colony network of roost-sharing between greater noctules in Maria Luisa Park, Seville, Spain.

Supplementary Figure S3 - Fig. S3 Network of parent-offspring dyads among giant noctule bats roosting in Maria Luisa Park, Seville, Spain.

## Data accessibility

Microsatellite genotypes, sample ID and location were deposited in the Dryad Digital Repository (doi: 10.5061/dryad.rc504). All data analysis scripts are accessible in the GitHub repository (doi: 10.5281/zenodo. 2653473).

## Conflicts of interest

None declared

## Notes

#### Summary of Updates

This version clarifies the hypotheses tested in this article. The scope of the article is narrowed to Nyctalus lasiopterus. Finally, a number of smaller corrections are also brought following an initial revision.

https://datadryad.org/resource/doi:10.5061/dryad.rc504/5

https://zenodo.org/record/2653473

## References

Aljanabi, S.M. & Martinez, I. (1997). Universal and rapid salt-extraction of high quality genomic DNA for PCR-based techniques. Nucleic Acids Res. 25, 4692–3.

Alcalde, J., Juste, J. & Paunović, M. (2016). Nyctalus lasiopterus. The IUCN Red List of Threatened Species 2016: e.T14918A22015318. http://dx.doi.org/10.2305/IUCN.UK.2016-2.RLTS.T14918A22015318.en. Downloaded on 24 November 2018.

Aric, A.H., Daniel, A.S., Pieter J. S. (2008). Exploring network structure, dynamics, and function using NetworkX. in Proceedings of the 7th Python in Science Conference (SciPy2008). Gäel, V., Travis, V., and Jarrod, M. (Eds). Pasadena, CA USA.

Barclay, R.M.R. & Harder, L.D. (2003). Life histories of bats: life in the slow lane. In Bat Ecology: 209–253. Kunz, T.H. & Fenton, M.B. (Eds.). University of Chicago Press.

Boston, E., Montgomery, I. & Prodöhl, P.A. (2008). Development and characterization of 11 polymorphic compound tetranucleotide microsatellite loci for the Leisler’s bat, *Nyctalus leisleri* (Vespertilionidae, Chiroptera). Conserv. Genet. 10, 1501–1504.

Burland, T.M. & Wilmer, J.W. (2001). Seeing in the dark: molecular approaches to the study of bat populations. Biol. Rev. 76, 389–409.

Carter, G.G. & Wilkinson, G.S. (2013). Food sharing in vampire bats: reciprocal help predicts donations more than relatedness or harassment. Proc. R. Soc. B Biol. Sci. 280, 2012–2573.

Castella, V., Ruedi, M. & Excoffier, L. (2008). Contrasted patterns of mitochondrial and nuclear structure among nursery colonies of the bat *Myotis myotis*. J. Evol. Biol. 14, 708–720.

Chaverri, G., Gillam, E.H. & Vonhof, M.J. (2010). Social calls used by a leaf-roosting bat to signal location. Biol. Lett. 6, 441–444.

Clutton-Brock, T. (2009). Cooperation between non-kin in animal societies. Nature 462, 51–57.

Clutton-Brock, T. & Lukas, D. (2012). The evolution of social philopatry and dispersal in female mammals. Mol. Ecol. 21, 472–492.

Dombrovski, V., Fenchuk, V. & Zhurauliou, D. (2016). New occurrence and the first breeding record of *Nyctalus lasiopterus* in Belarus. Vespertilio, 18: 55–59.

Dondini, G. & Vergari, S. (2000). Carnivory in the greater noctule bat (*Nyctalus lasiopterus*) in Italy. J. Zool. 251, 233–236.

Dubourg-Savage, M.J., Bec, J. & Gaches, L. (2013). First roosts of *Nyctalus lasiopterus* breeding females in France. Barbastella 6, 44–50.

Dray, S. & Dufour A. (2007). The ade4 Package: Implementing the Duality Diagram for Ecologists. J. Stat. Softw. 22, 1–20. doi: 10.18637/jss.v022.i04.

Emlen, S.T. & Oring, L.W. (1977). Ecology, sexual selection, and the evolution of mating systems. Science 197, 215–223.

Estók, P. & Gombkötő, P. (2007). Review of the Hungarian data of *Nyctalus lasiopterus* (Schreber, 1780). Fol. Hist. Nat. Mus. Matraensis 31, 167–172.

Fortuna, M.A., Popa-Lisseanu, A.G., Ibáñez, C. & Bascompte, J. (2009). The roosting spatial network of a bird□predator bat. Ecology 90, 934–944.

Gannon WL, Sikes RS. Guidelines of the American Society of Mammalogists for the Use of Wild Mammals in Research. J. Mamm. (2007). 88, 809–823. doi:10.1644/06-MAMM-F-185R1.1

Goudet, J. (1995). FSTAT (version 1.2): a computer program to calculate F-statistics. J. Hered. 86, 485–486.

Greenwood, P.J. (1980). Mating systems, philopatry and dispersal in birds and mammals. Anim. Behav. 28, 1140–1162.

Hamilton, W.D. (1964). The genetical theory of kin selection. J. Theor. Biol. 7, 1–52.

Hernández-Brito, D., Carrete, M., Ibáñez, C., Juste, J. & Tella, J.L. (2018). Nest-site competition and killing by invasive parakeets cause the decline of a threatened bat population. R. Soc. Open Sci. 5, 172477.

Ibáñez, C., Juste, J., Garcia-Mudarra, J.L. & Agirre-Mendi, P.T. (2001). Bat predation on nocturnally migrating birds. Proc. Natl. Acad. Sci. 98, 9700–9702.

Ibáñez, C., Guillén, A. & Bogdanowicz, W. (2004). *Nyctalus lasiopterus* — Riesenabendsegler. In Handbuch der Säugetiere Europas: 695–715. Krapp, F. (Ed.). Wiebelsheim: Aula-Verlag.

Ibáñez, C., Guillén, A., Agirre-Mendi, P.T., Juste, J., Schreur, G., Cordero, A.I., Popa-Lisseanu, A.G. (2009). Sexual segregation in Iberian noctule bats. J. Mammal. 90, 235–243.

Juste, J., Bilgin, R., Muñoz, J. & Ibáñez, C. (2009). Mitochondrial DNA signatures at different spatial scales: from the effects of the Straits of Gibraltar to population structure in the meridional serotine bat (*Eptesicus isabellinus*). Heredity 103, 178–187.

Kalinowski, S.T., Wagner, A.P. & Taper, M.L. (2006). Ml-Relate: a computer program for maximum likelihood estimation of relatedness and relationship. Mol. Ecol. Notes 6, 576–579.

Kaňuch, P., Fornůsková, A., Bartonička, T. & Bryja, J. (2007). Multiplex panels of polymorphic microsatellite loci for two cryptic bat species of the genus *Pipistrellus*, developed by cross-species amplification within the family Vespertilionidae. Mol. Ecol. Notes 7, 871–873.

Kerth, G. (2008). Causes and consequences of sociality in bats. Bioscience 58, 737.

Kerth, G., Mayer, F. & König, B. (2000). Mitochondrial DNA (mtDNA) reveals that female Bechstein’s bats live in closed societies. Mol. Ecol. 9, 793–800.

Kerth G, Mayer F, Petit E. 2002. Extreme sex-biased dispersal in the communally breeding, nonmigratory Bechstein’s bat (Myotis bechsteinii). Mol. Ecol. 11:1491–1498. doi: 10.1046/j.1365-294X.2002.01528.x.

Kerth, G., Perony, N. & Schweitzer, F. (2011). Bats are able to maintain long-term social relationships despite the high fission-fusion dynamics of their groups. Proc. R. Soc. B Biol. Sci. 278, 2761–2767.

Kerth, G. & Reckardt, K. (2003). Information transfer about roosts in female Bechstein’s bats: an experimental field study. Proc. R. Soc. B Biol. Sci. 270, 511–515.

Kovalov, V., Hukov, V. & Rodenko, O. (2019). New record of *Nyctalus lasiopterus* (Schreber, 1780) in Ukraine with a new confirmation of carnivory. North-West. J. Zool. 15, 91–95.

Kozhurina, E.I. (1993). Social organization of a maternity group in the noctule bat, *Nyctalus noctula* (Chiroptera□: Vespertilionidae). Ethology 93, 89–104.

Krebs, C.J. (1989). Ecological methodology. Harper & Row New York.

Lewis, S.E. (1995). Roost fidelity of bats: a review. J. Mammal. 76, 481–496.

Metheny, J.D., Kalcounis-Rueppell, M.C., Willis, C.K.R., Kolar, K.A. & Brigham, R.M. (2007). Genetic relationships between roost-mates in a fission–fusion society of tree-roosting big brown bats (*Eptesicus fuscus*). Behav. Ecol. Sociobiol. 62, 1043–1051.

Moussy, C., Hosken, D.J., Mathews, F., Smith, G.C., Aegerter, J.N. & Bearhop, S. (2013). Migration and dispersal patterns of bats and their influence on genetic structure. Mamm. Rev. 43, 183–195.

Nado, L., Chromá, R. & Kaňuch, P. (2017). Structural, temporal and genetic properties of social groups in the short-lived migratory bat *Nyctalus leisleri*. Behaviour 154, 785–807.

O’Donnell, C.F.J. (2000). Cryptic local populations in a temperate rainforest bat *Chalinolobus tuberculatus* in New Zealand. Anim. Conserv. 3, 287–297.

Oliphant, T.E. (2007). Python for scientific computing. Comput. Sci. Eng. 9, 10–20.

Olivera-Hyde, M.O., Silvis, A., Hallerman, E.M., Ford, M. & Britzke, E. (2019). Relatedness within and among *Myotis septentrionalis* colonies at a local scale. Can. J. Zool. 97, 724–735.

Patriquin, K.J., Palstra, F., Leonard, M.L. & Broders, H.G. (2013). Female northern myotis (*Myotis septentrionalis*) that roost together are related. Behav. Ecol. 24, 949–954.

Patriquin, K.J., Leonard, M.L., Broders, H.G., Ford, W.M., Britzke, E.R. & Silvis, A. (2016). Weather as a proximate explanation for fission–fusion dynamics in female northern longeared bats. Anim. Behav. 122, 47–57.

Popa-Lisseanu, A.G., Bontadina, F., Mora, O. & Ibáñez, C. (2008). Highly structured fission– fusion societies in an aerial-hawking, carnivorous bat. Anim. Behav. 75, 471–482.

Popa-Lisseanu, A. G., Bontadina, F. & Ibáñez, C. (2009). Giant noctule bats face conflicting constraints between roosting and foraging in a fragmented and heterogeneous landscape. J. Zool. 278, 126–133.

Popa-Lisseanu, A. G., Delgado-Huertas, A., Forero, M. G., Rodríguez, A., Arlettaz, R. & Ibáñez, C. (2007). Bats’ conquest of a formidable foraging niche: the myriads of nocturnally migrating songbirds. PLoS One 2, e205.

Rossiter, S., Jones, G., Ransome, R. & Barratt, E. (2002). Relatedness structure and kin-biased foraging in the greater horseshoe bat (*Rhinolophus ferrumequinum*). Behav. Ecol. Sociobiol. 51, 510–518.

Santos, J.D., Meyer, C.F.J., Ibáñez, C., Popa-Lisseanu, A.G. & Juste, J. (2016). Dispersal and group formation dynamics in a rare and endangered temperate forest bat (*Nyctalus lasiopterus*, Chiroptera: Vespertilionidae). Ecol. Evol. 6, 8193–8204.

Schuelke, M. (2000). An economic method for the fluorescent labeling of PCR fragments. Nat. Biotechnol. 18, 233.

Uhrin, M., Kaňuch, P., Benda, P., Hapl, E., Verbeek, H. J., Krištín, A., Krištofík, J., Mašán, P. & Andreas, M. (2006). On the greater noctule (*Nyctalus lasiopterus*) in central Slovakia. Vespertilio 9, 183–192.

Vlaschenko, A., Kravchenko, K., Prylutska, A., Ivancheva, E., Sitnikova, E. & Mishin, A. (2016). Structure of summer bat assemblages in forests in European Russia. Turkish J. Zool. 40, 876–893.

Vonhof, M.J., Davis, C.S., Fenton, M.B. & Strobeck, C. (2002). Characterization of dinucleotide microsatellite loci in big brown bats (*Eptesicus fuscus*), and their use in other North American vespertilionid bats. Mol. Ecol. Notes 2, 167–169.

Vonhof MJ, Strobeck C, Fenton MB. 2008. Genetic Variation and Population Structure in Big Brown Bats (Eptesicus fuscus): Is Female Dispersal Important? J. Mammal. 89:1411–1420. doi: 10.1644/08-MAMM-S-062.1.

Wagner, A.P., Creel, S. & Kalinowski, S.T. (2006). Estimating relatedness and relationships using microsatellite loci with null alleles. Heredity 97, 336–45.

Wilkinson, G.S. (1986). Social grooming in the common vampire bat, *Desmodus rotundus*. Anim. Behav. 34, 1880–1889.

Wilkinson, G.S. (1992). Communal nursing in the evening bat, *Nycticeius humeralis*. Behav. Ecol. Sociobiol. 31, 225–235.

Wilkinson, G.S. & Boughman, J.W. (1998). Social calls coordinate foraging in greater spearnosed bats. Anim. Behav. 55, 337–350.

Wilkinson, G.S., Carter, G., Bohn, K.M., Caspers, B., Chaverri, G., Farine, D., Günther, L., Kerth, G., Knörnschild, M., Mayer, F., Nagy, M., Ortega, J. & Patriquin, K. (2019). Kinship, association, and social complexity in bats. Behav. Ecol. Sociobiol. 73, 7.

Willis, C.K.R. & Brigham, R.M. (2004). Roost switching, roost sharing and social cohesion: forest-dwelling big brown bats, *Eptesicus fuscus*, conform to the fission–fusion model. Anim. Behav. 68, 495–505.

