## Supplementary Figure S3 for "Kin structure and roost fidelity in greater noctule bats"

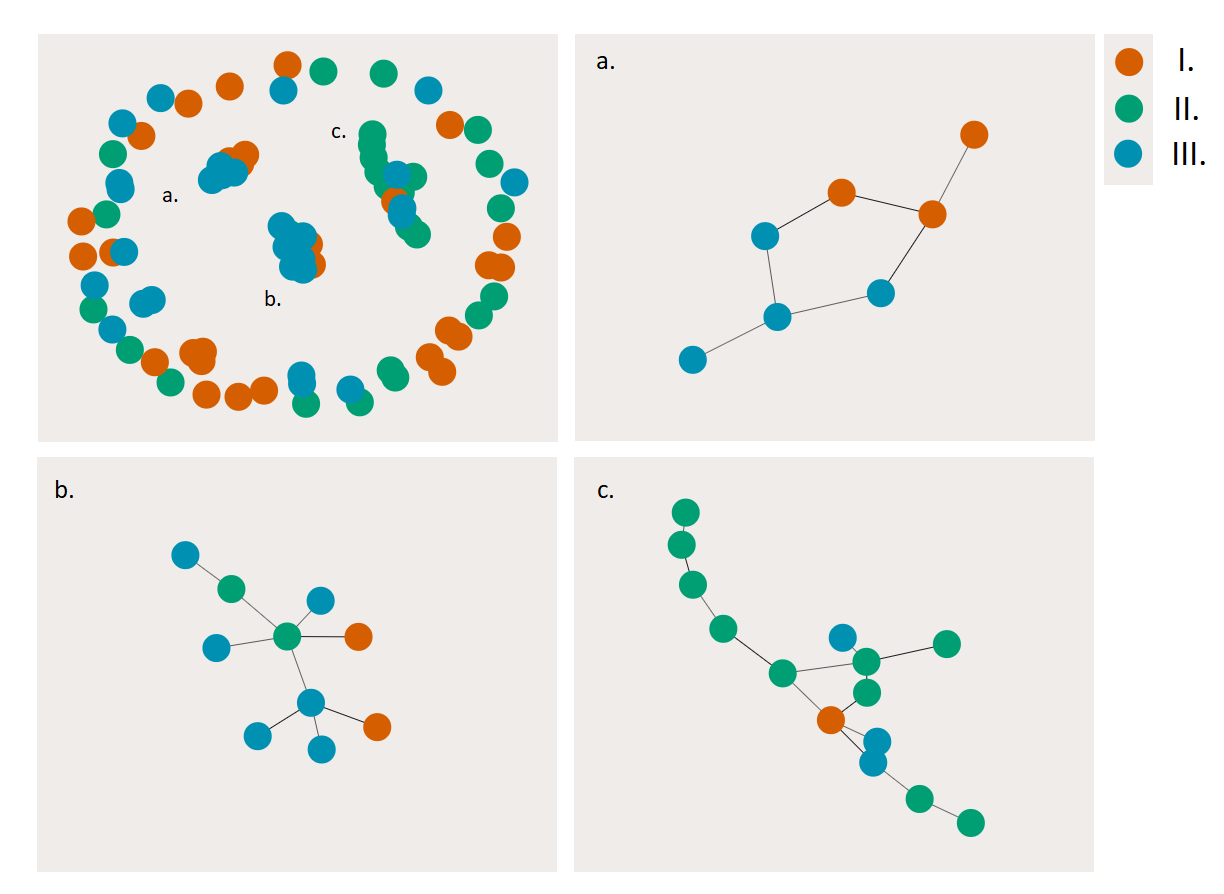

**Fig. S3 Network of parent-offspring dyads among giant noctule bats roosted in Park Maria Luisa.** The top-left panel displays all parent-offspring dyads within the Seville data set according to our analysis. Filial relations fall into three connected components. Each component is enlarged in the following panels. The network was constructed using the Fruchterman-Reingold algorithm. Strongly connected components enlarged. Colours represent assignment to social groups within the park (Popa-Lisseanu et al. 2008).
