## Supplementary Methods for "Kin structure and roost fidelity in greater noctule bats"

Supporting Information

*Detailed description of DNA extraction, purification, sequencing and genotyping*

Wing biopsy punches were first dipped into 300 µl of lysis buffer I (0.1 M Tris-HCL pH 8.0, 0.1 M EDTA pH 8.0) and 300 µl of lysis buffer II (0.1 M Tris-HCl pH 8.0 SDS 1%). 32 µl of NaCl 5M (0.28 M final concentration) and 30 µl of 20mg/ml proteinase K (873 µg/mll final concentration) were added, together with 25 µl of DTT. The samples were incubated at 55-60ºC overnight, after which 160 µl of NaCl 5M was added to each. Samples were vortexed for 10 min at 13 000 rpm, 4ºC for 10 minutes. The supernatant was transferred to fresh tubes, and an equal amount of isopropanol (~600 µl) was added. Samples were mixed, and incubated at -20ºC for 10 minutes, after which they were centrifuged for 10 minutes (13 000 rpm, 4ºC), the isopropanol removed, and each tube washed with 500 µl of ethanol 70%, dried, and then resuspended in 60-80 µl sterile dH_2_O. The DNA yield was 5 to 20 ng per sample, but the nanodrop readings should be regarded as highly skewed due to the salt levels in the final samples.

Amplifications of mitochondrial markers were conducted in 21 µl reaction volumes containing 3 µl of template DNA, 2 µl of buffer solution, 0.8 µl MgCl2, 0.8 µl of DMSO, 0.08 nM of dNTP, and 0.6 U of Taq DNA Polymerase (Quiagen). HVI reactions included 0.25 µM of each primer, and HVII reactions 0.38 µM. Thermal profiles were as follows: initial denaturation at 94ºC for 4 minutes, followed by 35 cycles consisting of 45 s at 94ºC, 45 s at annealing temperature (62ºC and 52ºC for HVI and HVII, respectively), and 1 min at 72 ºC.

In the case of mitochondrial fragments, PCR products were purified using Antarctic phosphatase buffer (BioLabs Inc., New England), and sequenced in the forward direction using a BigDye reaction mix (Applied Biosystems). The protocol for this step was the following: 2 µl of BigDye reaction mix, 2 µl of RNX Buffer, and 1µl of forward primer were added to the cleaned PCR products, resulting in a final volume of 12 µl. The sequencing products were cleaned of salts and unincorporated nucleotides by being filtered through Sephadex in 96-well plates, and read in a GS Junior (Roche 454) Sequencer.

Microsatellite genotyping was done following the procedure described by Schuelke (2000). Microsatellites were amplified independently, and the PCR products mixed according to the fluorescent labels used and the size range of the given markers. The 19 bp M13 tail sequence is added to the 5’ end of one of the primers (for simplicity, we modified only the forward primers). This sequence is identical to one attached to a fluorescent label added during the PCR reaction. The goal is for the latter to anneal to the former, functioning as a forward primer and producing labelled products.

To make this possible, 8 cycles are added at the end of the PCR. These retain the 94ºC and 72ºC denaturation and elongation steps respectively, but have an annealing temperature step of 53ºC. This temperature allows the M13 tail attached to the fluorescent label to attach itself to what is now, due to replication during previous cycles, a complementary M13 tail and forward primer. During these 8 cycles, due to the exponential increase of replicates during the initial cycles, it is possible to obtain a clear labelling of the amplified marker. The PCR conditions used were the following: 5 ng DNA, 1.4 to 2 mM MgCl2 (depending on the marker, see Table S1), 0.6 U of Taq Polymerase, 0.08 dNTP’s, 0.5 µl of non-modified reverse primer, 0.4 to 0.45 µM of M13 tailed forward primer, and 0.1 to 0.045 µM of labelled M13 tails.

The success of the labelling process depends on the proportion of M13 tailed primer to fluorescent labelled M13 tail. Personal communication with experienced researchers in the area lead to believe that an M13 label to M13_F primer ratio of 1:10 should be sufficient. Because Schuelke (2000) proposes the use of these in a 1:4 ratio we decided to test if there were significant differences of output using the different ratios and if so, which one gave the best results. We did this by amplifying 4 DNA extracts for the 11 microsatellites following both recipes and comparing the intensities obtained (height of the peaks when read). This was done for each marker separately. If the difference was significant (one-way ANOVA, P<0.05) we kept the recipe that produced the highest average peaks, if not, we used the 1:10 ratio. Results varied across markers, not always in the same direction. Because PCR reactions were performed separately, this did not make a difference though, and we adopted different recipes for different markers.
