## Supplementary Table S1 for "Kin structure and roost fidelity in greater noctule bats"

**Table S1** **Summary statistics and PCR specifications for each locus over all colonies** **the entire colony in Maria Luisa Park, Seville, Spain.** Includes annealing temperature (***T_a_***), total number of alleles (***A***), observed and expected heterozygosities (***H_o_*** and ***H_e_***), result of significance tests on deviations from Hardy-Weinberg equilibrium and estimates of null allele frequency (***F_null_***) at each locus.

| **Locus** | **Ta** | **MgCl_2_ (mM)** | **Allele size range (bp)** | ***A*** | ***Ho*** | ***He*** | ***Ar*** | ***HW^1^*** | **F(Null)** |
| --- | --- | --- | --- | --- | --- | --- | --- | --- | --- |
| *Nle3* | 50 | 2 | 219-251 | 93 | 0.667 | 0.761 | 5.7 ± 0.9 | NS | 0.0589 |
| *Nle6* | 58 | 1.4 | 213-276 | 92 | 0.88 | 0.81 | 10.9 ± 1.2 | * | -0.0636 |
| *EF4* | 54.5 | 1.4 | 238-252 | 89 | 0.787 | 0.786 | 5.6 ± 0.5 | NS | -0.0043 |
| *Nle10* | 58 | 2 | 149-165 | 89 | 0.427 | 0.509 | 4.6 ± 0.5 | NS | 0.0786 |
| *Nle11* | 50 | 2 | 158-188 | 86 | 0.744 | 0.863 | 8.4 ± 1.3 | ND | 0.0748 |
| *Nle2* | 52 | 2 | 203-233 | 91 | 0.813 | 0.818 | 9.2 ± 0.8 | NS | -0.001 |
| *Nle7* | 52 | 2 | 111-141 | 91 | 0.846 | 0.856 | 7.6 ± 0.6 | ND | 0.0022 |
| *Nle8* | 58 | 1.4 | 172-200 | 91 | 0.56 | 0.602 | 4.3 ± 0.7 | NS | 0.0353 |
| *Nle9* | 53 | 2 | 213-245 | 91 | 0.637 | 0.799 | 6.6 ± 1.3 | NS | 0.1105 |
| *P217* | 50 | 1.4 | 232-248 | 93 | 0.376 | 0.764 | 6.2 ± 0.4 | *** | 0.3331 |
| *P20* | 45 | 2 | 157-191 | 58 | 0.328 | 0.687 | 8.7 ± 1.7 | *** | 0.354 |

^1^NS: non-significant, **P<0.01, ***P<0.001
