## Supplementary Figure S1 for "Kin structure and roost fidelity in greater noctule bats"

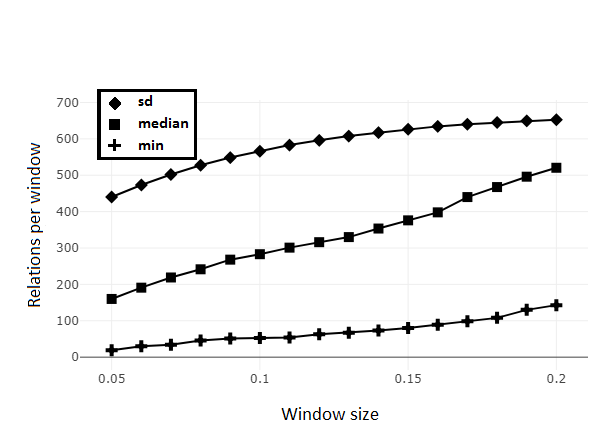

**Supplemental Fig. S1 Exploratory analysis of the impact of window size on sampling across relatedness steps**. Relatedness window size was varied between 0.05 and 0.2 at increments of 0.01. Sampling size for all windows of a given length overlapping at steps 0.01 was estimated.
