## Supplementary Figure S2 for "Kin structure and roost fidelity in greater noctule bats"

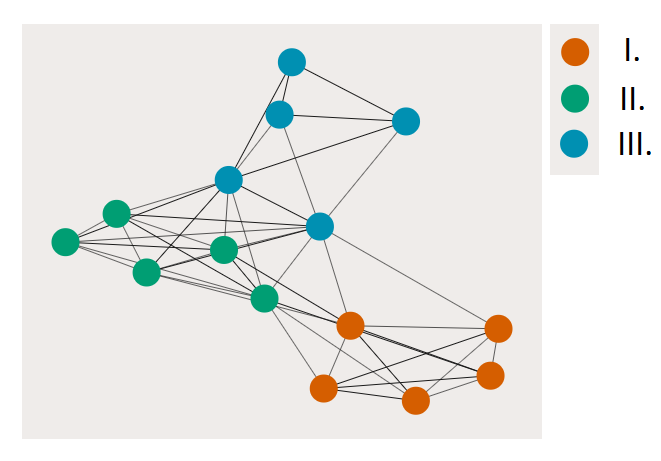

**Fig. S2 Intra colony network of roost-sharing between giant noctules in park Maria Luisa, Seville.** Nodes represent individual female bats. Edges were created as dyads presenting a similarity value above 0.05 (Freeman-Tukey statistic). Network was constructed using the Fruchterman-Reingold algorithm. Colours represent assignment to social groups within the park (Popa-Lisseanu et al. 2008).
